# Imaging and quantifying cytoplasmic tail dynamics of membrane receptors in living cells

**DOI:** 10.1101/2025.07.17.665247

**Authors:** Zhao Lin, Xinyao Chen, Lijuan Ma, Hang Fu, Hao Wang, Kai Yang, Ying Lu, Shuxin Hu, Ming Li

## Abstract

The cytoplasmic tails (C-tails) of membrane receptors are fundamental hubs for signal transductions. Yet directly investigating their dynamics in living cells is challenging. We developed a single-molecule technique to sense C-tail’s conformational transitions. It was applied to investigate the ligand-induced activation of β2AR, a prototypical G protein-coupled receptor (GPCR). Our data elucidate how the receptor transduces ligand binding to cell-level outcomes, with the ligand efficacy being interpreted in terms of the association and disassociation rates of the G protein. Moreover, upon G protein binding, not only the C-tail is displaced from the intracellular surface of the receptor but also the eighth helix H8 is dissociated from the plasma membrane. This work establishes a robust framework to directly decipher the activation of membrane receptors without interferences from overexpressed downstream transducers.

---

Conformational dynamics of membrane receptors are crucial for understanding the mechanisms of signal transductions. ^1-2^ This is particularly true for the cytoplasmic tails (C-tails) of membrane receptors, which are often intrinsically disordered^3-5^ yet serve as primary interfaces for intracellular signal complexes.^6-7^ Directly monitoring the C-tails in living cells is a formidable challenge. Addressing the challenge demands innovative experimental approaches and rigorous analytical frameworks to accurately interpret the results. The fluorescence resonance energy transfer (FRET) and the bioluminescence resonance energy transfer (BRET) are preferential for quantifying the protein conformational dynamics.^8-10^ However, they typically require two engineered fluorophores in one protein, which is usually not easy to realize in living cells. To date, single-molecule FRET/BRET has not yet been successfully used to study the C-tail dynamics of membrane receptors in living cells. In addition, the C-tails must interact with downstream proteins to transduce the signal. The ability to measure the tail dynamics with a single fluorophore may save a channel to visualize the receptor–transducer interactions if the effector protein is fluorescently labeled. In this work, we take advantage of the FRET from a single fluorophore to an ensemble of lipophilic dark quenchers being integrated into the cell membrane (Figure 1a). The approach has the very advantage of detecting the C-tail with only one fluorescent label. In addition, the sensitive detection range is from the inner surface of the plasma membrane to ∼10 nm underneath the plasma membrane, matching well the length of C-tails which are often a few nanometers long. We demonstrate the feasibility of our method by characterizing the dynamics of the C-tail of a prototypical GPCR, β2-adrenergic receptor (β2AR),^11^ which regulates numerous physiological processes including cardiac function, airway tone,metabolic function, and so on. We imaged directly the receptor’s basal activity, its suppression by an antagonist, and activation by a G protein-biased agonist. The single-molecule analysis revealed the conformational transitions of the C-tail and the reorientation of the eighth helix H8 of β2AR, which are both unseen in the structural biology but are strongly correlated with the G protein binding. Our approach enables quantitative assessment of the ligand efficacy and the receptor activity.

## RESULTS AND DISCUSSION

### Proof of principle and calibration of parameters

Our technique measures the FRET from a single fluorophore tagged to the C-tail to an ensemble of dark quenchers embedded in the membrane (Figure 1a). The fluorescence of the donor is attenuated by the dark quenchers diffusing in the membrane. The degree of the attenuation is characterized by F/F_0_ or τ/τ_0_, where F_0_ or τ_0_ is the intrinsic fluorescence intensity or the intrinsic fluorescence lifetime and the corresponding F or τ is measured after loading the quenchers. A standard curve of F/F_0_ or τ/τ_0_ as a function of distance (Figure 1b) can be numerically calculated using the basic energy transfer formulism (see Material and Methods in the Supplementary Materials for details), enabling determination of the donor-to-surface distance (denoted as z hereafter).

**Figure 1.**
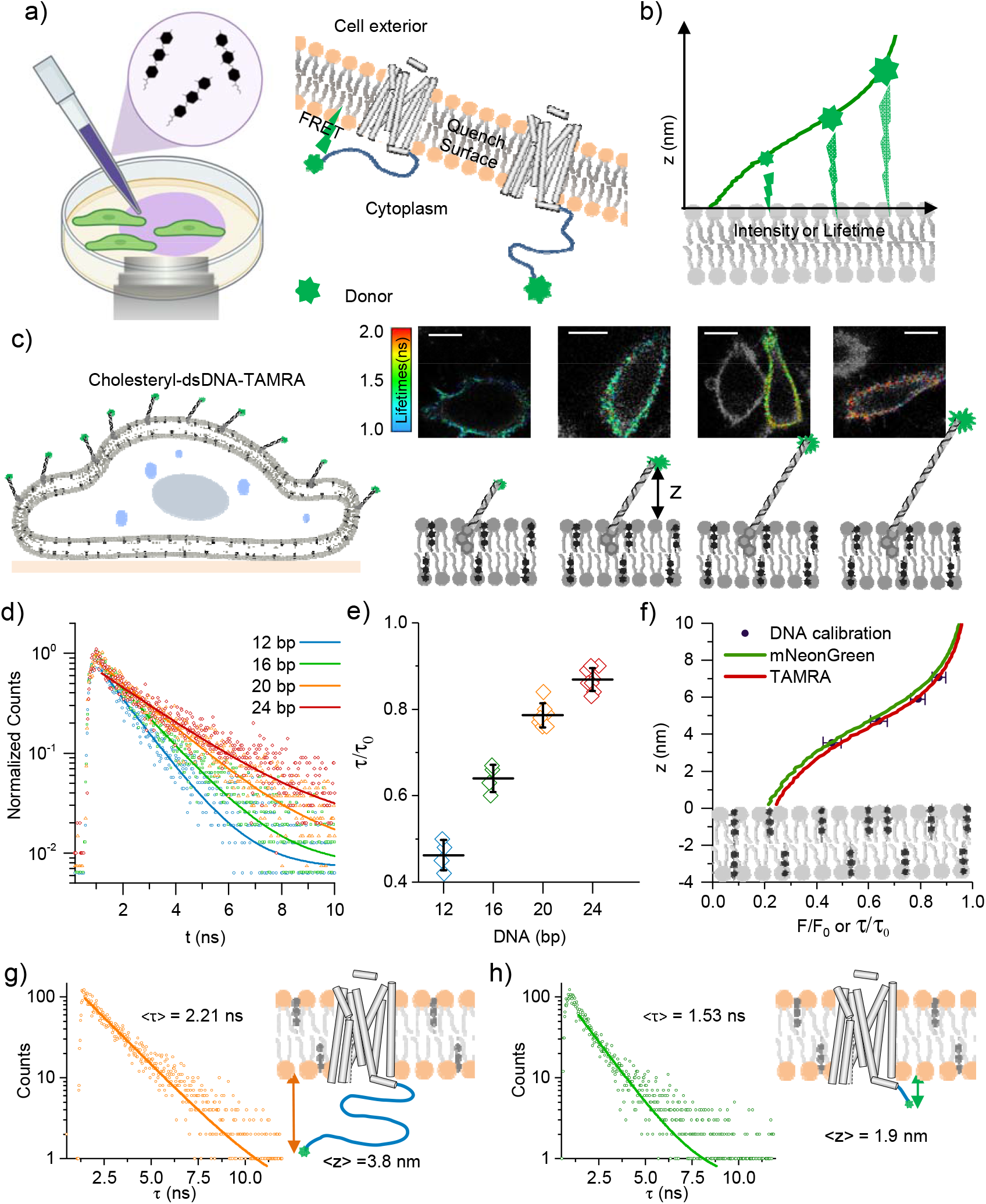
Proof of principle and calibration of parameters. a) Schematic diagram of sample preparation (left) and the principle of FRET from a single donor near the membrane to multiple quenchers inside the membrane (right). b) An expected relationship between F/F_0_ or τ/τ_0_ and the dye-to-surface distance z. c) Calibrating the 2D density of BHQ-1 by the FLIM images of cell surfaces with TAMRA-labeled dsDNA segments of various lengths in the presence of 15 μM BHQ-1. d) Histograms of the photon arrival times (lifetime curves) as measured by TCSPC. e) The lifetime ratios τ/τ_0_ obtained from different cells containing dsDNA segment of various lengths. f) Determining the density of BHQ-1 by comparing the τ/τ_0_ – z curve (red line) calculated by the Monte Carlo simulation with the experimentally measured curve. Also shown is the τ/τ_0_ – z curve (green line) for mNG. Error bars are standard derivations. g) The lifetime curve reflects the average donor-to-membrane distance in cells expressing β2AR–mNG following incubation with BHQ-1. h) The corresponding results for β2AR△C–mNG.

In the experimental implementation, the baseline fluorescence intensity (F_0_) and lifetime (τ_0_) measurements must precede quencher loading. Because the density of the quenchers in the membrane may become >5000 molecules per square micrometers during the measurements, they should be extremely dark to void photon leakage to the signal channel which may make the single-molecule measurements unfeasible. The black hole quenchers ^12^ (BHQ-1, Biosearch Technologies) have optimal spectral overlap with the fluorescent proteins we used (Figure S1), high extinction coefficient, broad absorption spectrum, and negligible emission.^12-13^ The positions of the quenchers in the membrane can be determined by molecular dynamic (MD) simulations. It turned out that the BHQ-1 molecules wedged into the headgroup regions of both leaflets of a lipid bilayer within 1 μs (Figure S2a). Given the 2-dimensional density and position of the quenchers in the membrane, the attenuation profile can be generated via Monte Carlo simulations ^14^ (Figure S2b). The cell viability assays confirmed the noncytotoxic nature of BHQ-1, with normal growth observed even one day after exposing to the quenchers containing culture medium (Figure S3). The analysis can be complemented using both the total internal reflection fluorescence (TIRF) microscopy for single-molecule tracking and the fluorescence lifetime imaging microscopy (FLIM) for single-cell analysis (Figure S4).

Before the actual live-cell analysis, we demonstrated the feasibility of our method by measuring the dye to surface distances of a series of dye-labeled double-stranded DNA segments standing on the surfaces of giant unilamellar vesicles (GUV) (Figure S5). We prepared the DNA with various lengths, which were labeled with a fluorescent dye at the upper end of the DNA and can be attached to the lipid membrane through cholesterol at the bottom end (Materials and Methods in the Supplementary Materials and Table S1). We found that the lifetime ratio (τ/τ_0_) measured with FLIM increases with the increasing length of the DNA segments when the GUVs were incubated with a fixed concentration of BHQ-1 (Figure S5). The quenching efficiency can be tuned by adjusting the BHQ-1 concentration (Figure S6). Referring to the reported tilt angle of the short dsDNA segments on a surface (about 60°),^15^ we obtained the experimental data of τ/τ_0_ vs z (Figure S5d), from which we can estimate the density of BHQ-1 in the membrane *via* Monte Carlo simulations mentioned above. We then calibrated the density of the quenchers integrated into the plasma membranes of the cells of interest by measuring τ/τ_0_ at various length of dsDNA (Figure 1c-f). To check the lateral homogeneity of BHQ-1 in the plasma membrane, we perform single molecular experiment with the DNA segments and found this technique is also compatible with the fluorescence intensity acquisitions (Figure S7).

### Validation of the technique for living cell assays

GPCRs are targets for more than one third of approved drugs, establishing them as fundamental to pharmacological research and therapeutic development.^16-18^ Assessing the drug-induced activities of GPCRs in physiological conditions is challenging due to their dynamic and transient interactions.^19-21^ We fused mNeonGreen (mNG, a green fluorescent protein) ^22-23^ to the C-terminus of β2AR (β2AR–mNG) as the donor (Figure 1g). As a control, we engineered a truncated β2AR variant (β2AR△C - mNG) by removing the tail after Ser346 and connecting this residue to mNG *via* a 10-amino-acid linker (Figure 1h). Following 30-minute incubation with 15 μM BHQ-1, the cell-level FLIM measurements gave a lifetime ratio τ / τ_0_ ≈ 0.68 for wild-type β2AR, indicating an average C-terminus-to-surface distance of ∼3.8 nm. In contrast, the truncated variant exhibited a short distance of ∼1.9 nm (τ / τ_0_ ≈ 0.51). We also performed single-molecule measurements using TIRF (Figure 2 and Figure S8a). Both the intensity ratio F/F_0_ and the lifetime ratio τ/τ_0_ reveal that the C-terminus of the truncated variant resides closer to the membrane than that of the full-length receptor, demonstrating the feasibility of our technique in characterizing C-tails.

**Figure 2.**
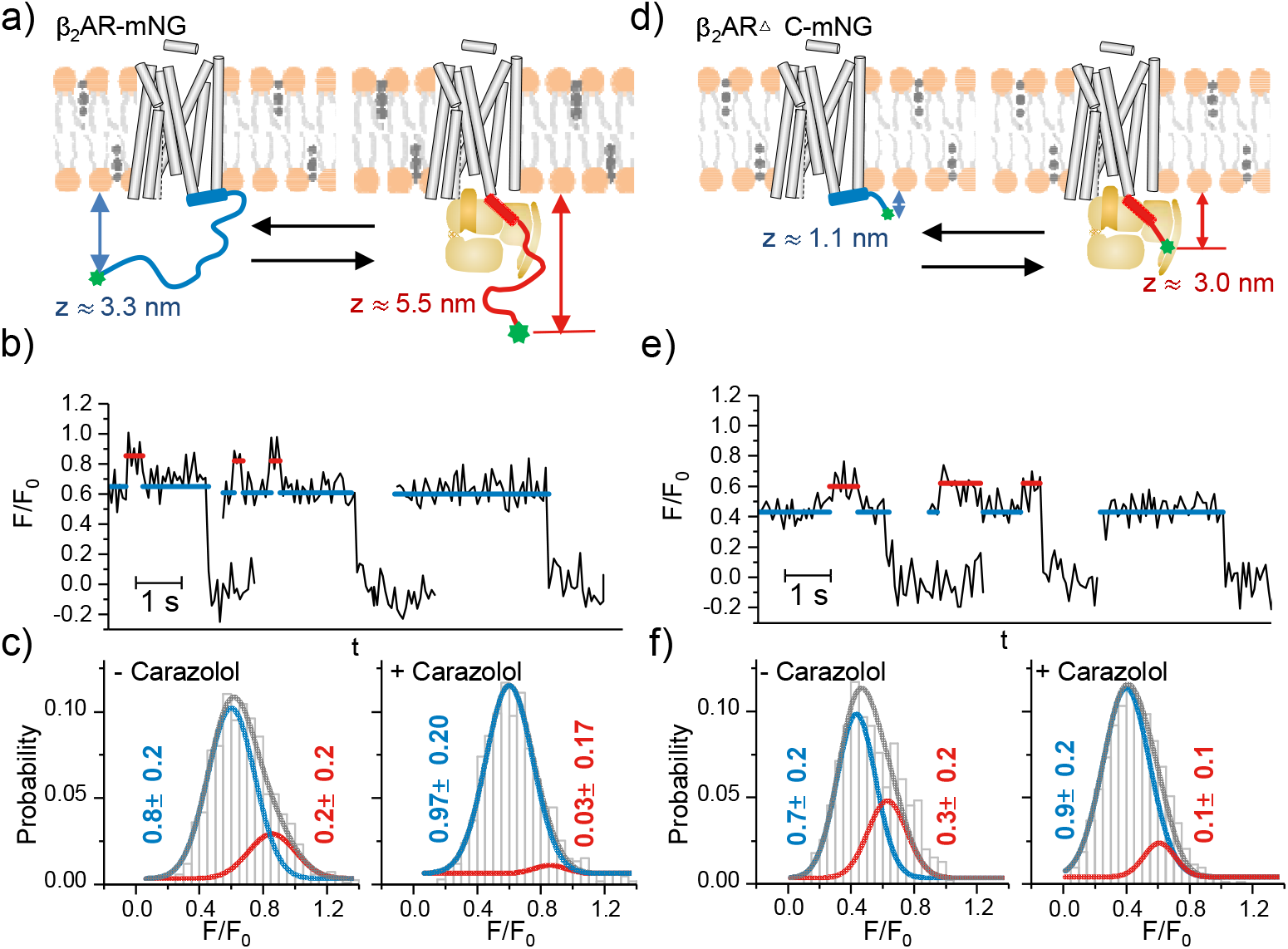
Single-molecule analysis of the C-tail in resting cells and in antagonist-inhibited cells. a) Schematic diagram of the transitions of the C-tail in β2AR–mNG between the compact (blue) and the extended (magenta) conformations. b) Representative time traces of F/F_0_ for β2AR–mNG. c) Histograms of the conformations, each being fitted to a combination of two Gaussians. The area of each Gaussian is given at the peak. d-f) The corresponding results for β2AR△C–mNG.

### The C-tail is in extended conformation upon G protein binding

We observed that, even in the absence of any stimulation, the single-molecule fluorescence traces displayed switches between a state with F/F_0_ ≈ 0.65 (z ≈ 3.3 nm; blue line) and a state with F/F_0_ ≈ 0.85 (z ≈ 5.5 nm; magenta line) (Figure 2a, b and Figure S8b). That is, the C-tail of the receptor samples at least two distinct conformations: a dominant compact one and an occasional extended one (Figure 2c). The compact conformation agrees with molecular-dynamics simulations showing that the electrostatic attraction between the negatively charged C-tail and the positively charged intracellular loop 3 can tether the tail close to the membrane.^24^ The origin of the extended conformation, however, is not obvious.

We assume that the extended conformation is due to the G protein binding to the activated GPCR for the following reasons: 1) There exists basal activation of GPCR in unstimulated cells which enables sporadic, low-level cAMP signaling.^25-27^ A previous in vitro study showed that G proteins may bind spontaneously to unstimulated GPCRs.^9,28^ Such binding is expected to disrupt the electrostatic interaction between the C-tail and the cytoplasmic surface of the receptor, releasing the tail from its membrane-proximal tether.^24^ 2) It is well known that the antagonist carazolol inhibits the G proteins from binding to β2AR.^29^ The addition of carazolol (10 μM) suppressed almost all the basal activation events (Figure 2b, c and Figure S8c). 3) As established, the G protein biased agonist ICL 3-9 promotes selectively G protein binding.^30,31^ When β2AR was stimulated by 10 μM ICL 3-9, the proportion of the extended conformation increased substantially (Figure 4a and Figure S8d). Altogether, the G protein binding could displace the C-tail from the GPCR’s intracellular surface so that it transits to the extended conformation. Therefore, the transient excursions between the compact and the extended conformations may arise from the binding and unbinding of the G proteins.

### The H8 helix is involved in the activation process

A simple worm-like-chain model estimates that the displaced C-tail would have an average extension of ∼4.5 nm (please see Supporting Information for details), which is notably shorter than the measured value of 5.5 nm. The discrepancy can be explained by the observation that the C-tail truncated receptor β2AR^△^C-mNG also exhibited two discrete conformations: a membrane-proximal one (F/F_0_ ≈ 0.43, z ≈ 1.1 nm; blue line in Figure 2e, f) and a membrane-distal one (F/F_0_ ≈ 0.60, z ≈ 3.0 nm magenta line). The 1.9 nm increment corresponds approximately to the length of the H8 helix of β2AR, suggesting a dramatic reorientation of the helix. The population of the membrane-distal conformation increased from ∼30% to 50% when the cells expressing β2AR△C-mNG were stimulated with the G protein biased agonist ICL 3-9 (Figure S9). It suggests that even in the absence of the C-tail, H8 itself undergoes basal conformational transitions between a state parallel to the membrane and a state perpendicular to the membrane. We therefore speculate that the observed 5.5 nm distance may reflect a combined displacement of both the C-tail and the H8 helix.

To further validate whether the movement of H8 is correlated with the G protein coupling, we engineered a β2AR mutant (β2AR^T68F,Y132G,Y219A^△C; β2AR^TYY^△C) that is incapable of activating G proteins.^32-33^ Indeed, in cells expressing β2AR^TYY^△C-mNG, the membrane-distal conformation was barely detectable under the basal condition (10% compared with 30% in β2AR△C-mNG), and stimulation with ICL 3-9 failed to increase its population significantly (Figure 3). It is noteworthy that the crystal structures of β2AR do not show large H8 movements upon activation,^34-35^ although biochemical evidences identify H8 as a critical site for the G protein engagement.^36-37^ Our *in vivo* results raise the possibility that the H8 reorientation might only become apparent when the GPCR is in native plasma membranes.

**Figure 3.**
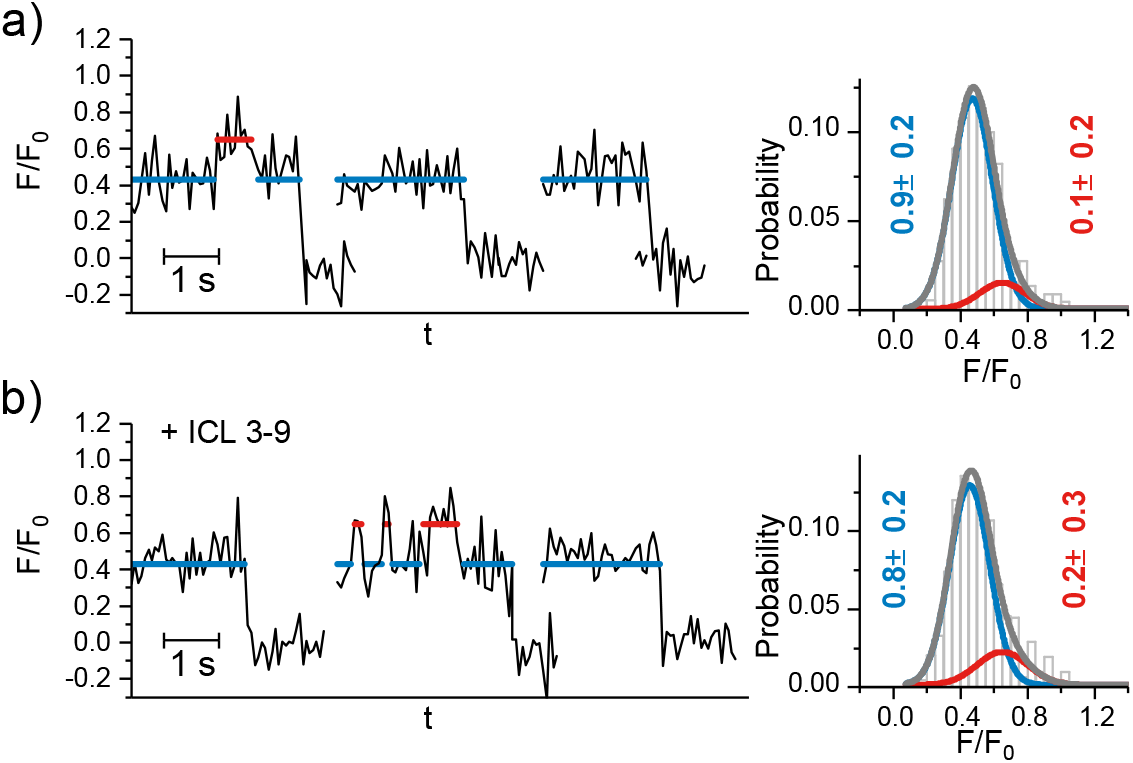
Single-molecule analysis of the β2AR^TYY^ C-mNG in resting cells and in ICL 3-9 stimulated cells. a) Representative time traces of F/F_0_ for β2AR^TYY^_△_C-mNG. The histograms of the conformations are displayed on the right. b) The corresponding results for β2AR^TYY^△C-mNG stimulated by ICL 3-9.

### The G protein binding kinetics of GPCR upon activation

ICL 3-9 promotes selectively G protein coupling without inducing GRK phosphorylation or β-arrestin recruitment.^30-31^ However, desensitization *via* alternative pathways such as PKA-mediated phosphorylation still occurs.^38^ We studied the activation of β2AR by ICL 3-9. Again, two states were observed (Figure 4a). Their relative proportions changed with time as can be seen from the representative fluorescence traces and the histograms in different time windows. ICL 3-9 led to an initial increase of the population of the extended conformation from ∼20% to ∼40% within 1 min, and then a slow restore to ∼20% after 20 min. The increase of the dwell time of the active state from 0.21 s in the basal condition to 0.34 s upon ICL 3-9 stimulation (Figure 4b) indicates ligand-induced stabilization of the G protein binding. Our results agree well with an in vitro result in the presence of GDP and GTP.^9^ Interestingly, we found that the dwell time declines to 0.28 s ultimately, suggesting that the desensitization of the GPCR only weakens, rather than blocks the G protein–GPCR interaction. We will show in the following that the dwell-time values provide an effective measure of the ligand efficacy in the cell level analysis.

**Figure 4.**
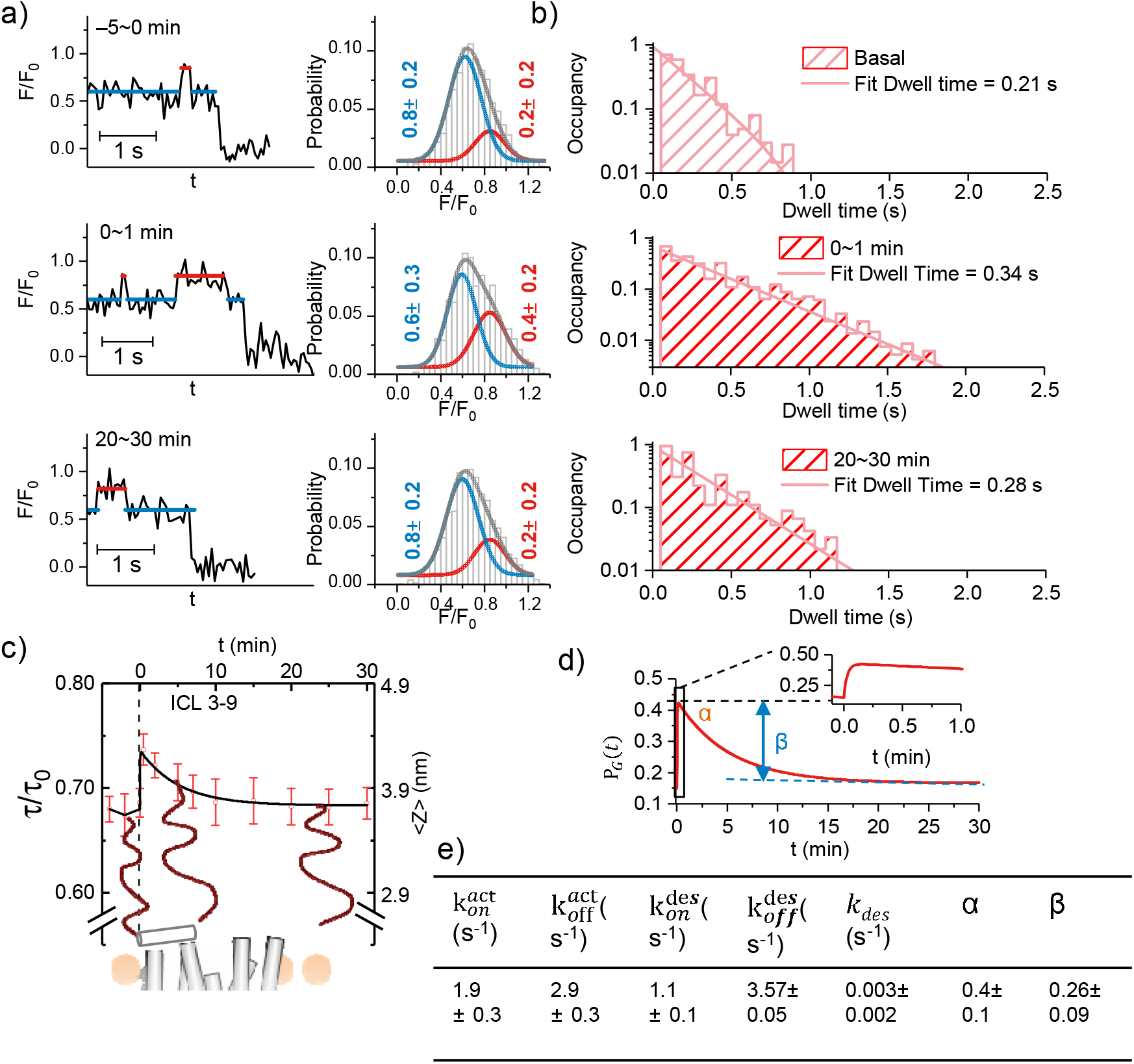
Conformational changes of the C-tail induced by the G protein-biased agonist. a) Representative intensity traces collected at different time windows after the ICL 3-9 loading. The histograms of the conformations are displayed on the right. b) Dwell times of the extended state in resting, immediately activated and desensitized cells. The statistics were from >100 events. c) Time course of the lifetime measured by time-lapse FLIM and the schematic of the corresponding C-terminus-to-surface distance (red cycles with error bars). The black line represents the theoretical curve derived from the kinetic model. d) The occupation probability P_G_(t) of a GPCR by a G protein using the kinetic parameters in e).

### The molecular-level dynamics is mirrored by the cellular signaling responses

Our results enable an analytical description of recruitment of G proteins in the native cellular context. To show this, we performed time-lapse FLIM images to further display the time evolution of the C-tail’s conformation (Figure 4c). ICL 3-9 led to a biphasic conformational change: a rapid initial extension followed by a gradual retraction. The pattern reflects a picture that rapid G protein recruitment is followed by slow receptor desensitization.^39^ Similar features were reported for many GPCRs but have not yet been understood in the molecular details. We propose an effective kinetic model to transform the single-molecule kinetics to the ligand efficacy. Assuming that all the agonists bind to the membrane receptors at t = 0, the ICL 3-9 stimulation enhances the G protein recruitment to rapidly increase the G protein-bound population, followed by a gradual decline of the population due to desensitization.^38-39^ To interpret our single-molecule and FLIM data within a unified framework, we constructed an effective kinetic model that coarse-grains the complex conformational landscape of the C-tail and the H8 helix into two signal-defined states: a G-protein-unbound state (with τ_*unbound*_/ τ_0_ ≈ 0.65) and a G-protein-bound state (with τ_G_/ τ_0_ ≈ 0.85). This reduction is justified because the measured donor-to-membrane distance is a projected observable of a higher-dimensional conformational distribution, as is common for intrinsically disordered regions and polymer-like tethers. The association and dissociation of G proteins to GPCRs and the GPCR activation-induced desensitization can be described by three kinetic equations (see Eq. S1, S3 and S5 in the Supporting Information), the solutions of which give the time evolution of the probability of a GPCR being bound by a G protein,

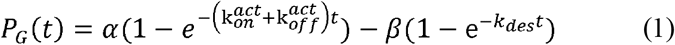

Where 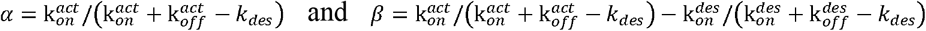. For the basal state prior to agonist stimulation (t < 0), the G protein binding was in a steady state, described by 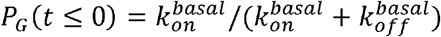. The parameters αand β can be considered as the apparent efficiencies of activation and desensitization, respectively. The parameters 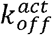 (dissociation rate of the G proteins from the activated receptors) and 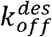 (dissociation rate of the G proteins from the desensitized receptors) can be estimated directly from the single-molecule dwell times shown in Figure 4b, under the assumption that the extended-state lifetimes report on the G-protein-bound duration. In principle, 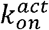 (association rate of the G proteins to the active receptors) and 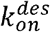 (association rate of the G proteins to the desensitized receptors) could also be directly extimated from the the single-molecule measurements if the traces were long enough. Unfortunately, we were not able to directly measure the durations of a G protein in the dissociate state due to photobleaching-induced shortening of the traces. Instead, the kinetic parameters 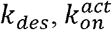 and 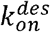 are inferred as effective parameters by globally fitting the time-resolved FLIM data (Figure 4c) to the ensemble-averaged prediction of the kinetic model (see Eq. S10):

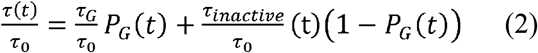

where the lifetime ratio of the G protein-unbound GPCRs is τ_*unbound*_/τ_0_ ≈ 0.65 and that of the G protein-bound GPCRs is τ_*G*_/τ_0_ ≈ 0.85, both were given by the single-molecule measurements. The experimentally measured cell-level time evolution of the lifetime ratio (Figure 4c, red cycles) decays very slowly, yielding an effective desensitization rate constant *k*_*des*_ = 0.003/s. As *k*_*des*_ is much smaller than the effective association-dissociation sum 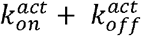 , justifying the approximation used to extract the activation parameters from the early time points, the maximum of the FLIM curve (Figure 4c, red point) can be used to approximate *p*_*G*_ (*t* = 30 s) ≈ *α* (see also the inset in Figure 4d). This gives *α* and 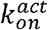 because 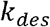 and 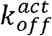 areknown. Similarly, *p*_*G*_(*t* = 1200 s) ≈ *α*-*β*. This gives *β* and then 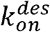. The resulting parameters are listed in Figure 4e, which reproduce very well the FLIM data we measured (Figure 4c).

We applied our tool to the full agonist isoproterenol (ISO) to further validate the quantitative capacity of our method. we used the C-tail truncated construct β2ARΔC-mNG to isolate the G-protein-dependent signaling. Comparing the full agonist isoproterenol (ISO) with the G-protein-biased agonist ICL 3-9, we found that ISO induced higher activation and desensitization efficacy (Figure S10).

## CONCLUSION

The C-tail of a membrane receptor serves as the direct indicator of receptor activation,^40-41^ which was often investigated using indirect biosensors involving transducers and/or second messengers that may introduce system cross-talks and lead to overestimation of the efficacy due to certain amplification mechanisms.^42-43^ Our approach migrates these shortages by directly monitoring the conformational dynamics of the C-tails. It captures not only the degree of activation of the receptor but also its temporal evolution. We demonstrated the powerfulness of our technique by using β2AR as a model. The observed conformational spectrum provides a structural basis for understanding the G protein-biased activation of β2AR. The established kinetic model transforms the ligand-induced changes in the G protein association and disassociation rates to the increment in proportion of the G protein-bound receptors, offering a new framework for ligand efficacy evaluation.

Remarkably, our work provides a new insight into the dynamics of the H8 helix in β2AR. We observed outward movement of the H8’s C-end upon activation—a dramatic conformational change contrasting with crystallographic studies which showed membrane-parallel H8 orientations in both the active and the inactive states. The dissociation of the amphiphilic helix from the plasma membrane may induce partial unfolding of the helix. This result agrees with a report that the agonist binding may cause H8 to lose part of its helical structure.^44-45^ Recent cell-level FRET studies on mechanosensitive GPCRs such as the histamine H1 and the B2 receptors revealed that mechanical stimulations may lead to increase of the distance between the H8 C-terminal and N terminal or between the H8 C-end and the cytoplasmic end of TM6. ^46-47^ Our data suggest that these changes may reflect complete H8 membrane dissociation rather than simple spring-like extension,^48^ potentially revising the current understanding of the H8 dynamics.

## EXPERIMENTAL SECTION

### Principle of the method

#### Distance dependence of the fluorescence signal

Our method is essentially the FRET between a single fluorophore near the plasma membrane and an ensemble of quenchers integrated in the membrane. The energy transfer kinetics k_t_ is the sum of pairwise transfer rates,

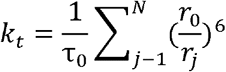

where τ_0_ is the intrinsic lifetime of the donor, r_0_ the F□ rster distance between the donor and a quencher molecule, and r_j_ the spatial distance between the donor and the j^th^ quencher. The FRET efficiency is

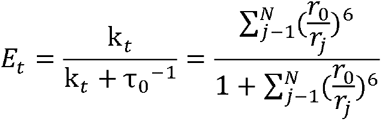

The experimentally measured relative fluorescent intensity F/F_0_ and/or the relative lifetime τ/τ_0_ are given by

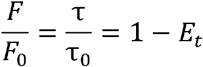

Assuming that the quenchers diffuse in the membrane, the energy transfer efficiency can be calculated by using the Monte Carlo simulations proposed in Ref. 14.^14^

#### Molecular dynamic (MD) simulation of the position of quenchers in the membrane

The spatial distance r_j_ between the donor and the j^th^ quencher depends on the position of the donor to be measured and the position of the quenchers that can be determined by MD simulation, which was performed by using the GROMACS 2021.4 package (www.gromacs.org) with the MARTINI force field.^49^ The simulation was carried out in the semi-isotropic ensemble at a temperature of 300 K and a pressure of 1.0 bar.^50-51^ The all-atom model of BHQ-1 was built by the ChemDraw software and coarse-grained using the method developed by Miller *et. al*..^52^ A bilayer consisting of 720 1,2-Dioleoyl-sn-glycero-3-phosphocholine (DOPC) lipids was constructed, 4 BHQ-1 molecules were added to both monolayers or the bilayer midplane. At least three independent runs were performed for at least 10 μs with a time step of 20 fs to ensure computational consistency.

### Sample preparations for imaging

#### Preparation of GUVs

The GUVs used in the calibration were generated *via* electro-formation using a Nanion Vesicle Prep Pro setup (Nanion Technologies), following the protocol provided in Ref. 53.^53^ The lipid compositions are 20% (by mol/mol) 1,2-dipalmitoyl-sn-glycero-3-phosphocholine (DPPC), 1,2-dioleoyl-sn-glycero-3-phosphocholine 10% (DOPC), 50% 1-palmitoyl-2-oleoyl-glycero-3-phosphocholine (POPC), 0.1% 1,2-dipalmitoyl-sn-glycero-3-phosphoethanolamine-N-(cap biotinyl) (Biotinyl-cap PE) and 20% cholesterol. The GUVs were added into a flow chamber with a streptavidin-coated glass surface^14^ and incubated for 2 min before each measurement.

#### The plasmids

The β2AR-mNeonGreen fusion construct was generated by PCR amplification of both genes. The β2AR sequence was amplified using a forward primer (5′-GCG CGC CAT GGG GTA CCC ATA CGA CGT CCC AGA CTA CGC CCA ACC CGG GAA CGG CA-3′) and a reverse primer (5′-TGG ATC CCG GGC CCG CGG TAC CGT CGA CCA GCA GTG AGT CAT TTG TAC TA-3′) containing BssHII and BamHI restriction sites. mNeonGreen was amplified with a forward primer (5′-GGA TCC ATC GTG AGC AAG GGC GAG GA-3′) incorporating a BamHI site and a reverse primer (5′-TCT AGA TCC GGA TCA CTT GTA CAG CTC GTC CAT-3′) with an XbaI site. The digested PCR products (BssHII/BamHI for β2AR; BamHI/XbaI for mNeonGreen) were ligated into the BssHII/XbaI-digested pEGFP vector (Clontech). A truncated β2AR variant (after S346) was created by substituting the reverse primer with 5′-GGA TCC CGG GCC CGC GGT ACC GTC GAC AGA AGA CCT GCG CAG G-3′. The G-protein-uncoupled truncated β2AR TYY mutant (T68F, Y132G, Y219A) was generated by site-directed mutation using the truncated β2AR variant as a template. All constructs were verified by DNA sequencing (BGI, Shenzhen).

#### The cells

Hela cells (ATCC, USA) were cultured in Dulbecco’s modified Eagle medium (DMEM, Corning) with 10% (vol/vol) fetal bovine serum (FBS, Gibco, Life Technologies, USA) and 1% penicillin-streptomycin (Gibco, Life Technologies, USA) medium maintained in a humidified incubator with 5% CO2 at 37 °C. The cells were routinely tested for mycoplasma. In general, when the cell confluency reached ∼80%, the cells were treated with 0.25% trypsin (Gibco, Life Technologies, USA) and passaged or transfected. Transfection was performed with Lipofectamine 3000 (Invitrogen, USA) according to the manufacturer’s instructions. Five hours after transfection, the cells were digested with trypsin and plated onto 35 mm confocal dishes with glass bottoms (Cellvis, USA) and cultured for another 6-24 h before imaging. Before imaging, medium was changed to live cell imaging buffer (Invitrogen, USA).

#### The fluorophore-labeled DNA

All DNAs were purchased from Sangon Biotec, China. One of the ssDNA was labeled with tetramethylrhodamine (TAMRA) at the 5’-end. The complementary ssDNA was labeled with cholesterol at the 5’-end. The dsDNA was diluted to 100 nM for the FLIM measurements and 0.1 nM for the TRIF measurements.

### The imaging and calibration methods

#### The calibration process using GUVs

The 2-dimensional density of the quenchers in the membrane should be determined at first to calculate the distance dependence of the fluorescence signal by using the Monte Carlo simulations. To this end, we calibrated the FRET efficiency at various positions by measuring the lifetimes of fluorophores attached to a series of dsDNA segments with lengths 12, 16, 20, 24 and 28 bp (Table S1 and Figure S5). The vesicles were immobilized on the bottom of the sample chamber through biotin–streptavidin interaction.^14^ The DNA samples were anchored by the cholesterol molecules to the surface of the lipid vesicles. The intrinsic fluorescence lifetime τ_0_. Was measured at first. Then the buffer with BHQ-1 was flowed in. The lifetime τ was measured after incubation for 30 min.

#### The live cell imaging

Before the imaging, the cells were transferred into the live-cell imaging buffer (Thermo Fisher Scientific) to maintain the physiological conditions. We first calibrated the density of BHQ-1 in the plasma membrane following the procedure used in the vesicle experiments (Figure 1). It is worth noting that the lipid vesicles have higher BHQ-1 adsorption than the live cells. We measured the cells transfected with fluorophore-labeled proteins. After measuring the τ_0_ and τ (or F_0_ and F), the quenchers were added to the living cell imaging solution (from Thermos Fisher) and incubated for 30 min before each measurement. We then introduced the agonist and monitored the fluorescence changes with time. The cells of interest were imaged with FLIM and/or with TIRF.

The TIRF measurements were performed on an objective-based TIRF microscope (IX71, Olympus) at room temperature. An oil-immersion objective (100x, NA 1.49) was used. The fluorescence signals from the cell were collected by an EMCCD (iXon, Andor Technology) with a 50 ms exposure time per frame. The fluorescence molecules were excited by a 488 nm Sapphire laser (Coherent Inc.). The imaging process was controlled and recorded by MetaMorph software (Molecular Device, California). The FLIM measurements were performed on a confocal laser scanning microscope (FV3000, Olympus) equipped with a multi-channel picosecond event timer (Hydraharp 400, PicoQuant) with 16 ps time resolution. The excitation unit consisted of a pulsed diode laser (λ_exc_ = 488 nm, LDH-D-TA 488, 20 mW, PicoQuant) with a pulse width 70 ps FWHM and a repetition rate 80 MHz. A high numerical aperture objective (UPLSAPO 100x oil, NA 1.49, Olympus) was used to focus the light into the sample and collect the fluorescence emission.

### Theoretical estimation of the extension of the disordered C-tail

To estimate the expected extension of the C-tail being displaced from the intracellular surface of the GPCR, we modeled it as a flexible, unstructured polypeptide chain in a good solvent. The contour length L_c_ of the C-tail, not including H8, is the production of the number of the residues (N = 82) multiplied by the length per residue (l = 0.38 nm), yielding L_c_ ≈ 31.16 nm. The mean squared end-to-end distance for a free chain is 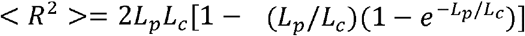, where the persistence length L_p_ was taken to be 0.8 nm for an unstructured polypeptide under physiological conditions. This yields a value of R ≈ 7 nm.

In our experimental geometry, the tail is anchored at the membrane surface at one end, imposing a constraint that the chain must remain in the half-space underneath the membrane. What we measured was the end-to-distance, < *z* >, of the chain. We compute the mean of the vertical component z over the accessible half-space. For a chain whose free end follows a spherically symmetric distribution *P*_0_(*z*) in the absence of a wall, the restricted distribution in the half-space *z*>0 is P(z) = 2*P*_0_(*z*) The average vertical distance is therefore

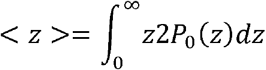

Using a Gaussian approximation for P_0_(z) as the contour length is much longer than the persistence length, this integral results in a value of < *z* > ≈ 4.5 nm nm. This is the maximum value one can expect. Any deviation from the free chain model, such as the folding into a local helix folding or binding to the surface of the G protein, would make it smaller. However, the value is notably smaller than we measured in the main text.

### The kinetics of transducers after addition of the ligands

#### Effective coarse-grained description

The following kinetic equations are not intended as a microscopic state model that maps one-to-one onto discrete structural conformations of the receptor–G-protein complex. Rather, they provide an effective coarse-grained description of the signal-defined dynamics observed in our experiments.^54,55^ The C-tail of β2AR is intrinsically disordered, and the measured donor-to-membrane distance represents a projected average over a distribution of tail conformations, H8 orientations, and possible membrane-proximal interactions. The two states we identify — with τ/ τ_0_ ≈ 0.65 and 0.85 — are therefore operational states defined by the fluorescence readout, not necessarily single structural species. Moreover, G-protein binding and dissociation occur on a crowded membrane surface where local rebinding and incomplete spatial mixing may renormalize apparent rate constants. Hence, the parameters derived from this model should be interpreted as effective kinetic coefficients that capture the dominant timescales of the signaling dynamics under our experimental conditions, rather than as microscopic rate constants in dilute solution.

Let N_0_ be the total number of GPCRs on the cell membrane. Because the agonist was applied at a saturating concentration of 10 μM, far exceeding the receptor numbers on the cell. Although the exact dissociation constant K*_d_* for ICL 3-9 has not been reported, it is likely in the submicromolar to low micromolar range based on the functional EC_50_ values (∼1.5 µM for ICL3-9). Taking a conservative estimate of K*_d_* ≈ 0.1∼1 µM, the applied concentration (10 µM) is at least 10-fold higher, ensuring that the receptor is saturated by the agonist and the binding equilibration time is estimated to be < 0.1 s. As binding relaxation time can be estimated as:

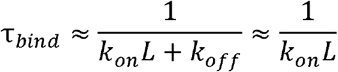

Where k_on_ is the diffusion-limited association rate constant ∼ 10^6^ − 10^8^ M^-1^s^-1^ for small peptides to GPCR, and k_off_ is negligible in the saturating regime. Taking L = 10 µM in our experiments, we can obtain τ_bind_ < 0.1*s*, which is much small than the temporal resolution of our FLIM measurements (≈ 1 min per frame). So we approximate that the agonist binding is instantaneous and that all receptors are in the agonist-bound state at t = 0. Because the rate of desensitization, including the one mediated by PKA, is much slower than the rate of G protein recruitment, the former can be considered independently of the latter. The change with time of the number of desensitized GPCRs N_des_ (t) follows

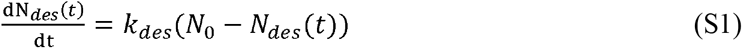

where *k_des_* is the effective desensitization rate. The solution of Eq. (S1) is

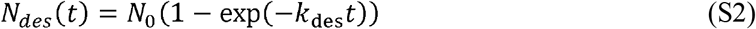

Assuming that the desensitized GPCRs recruit the G proteins in the rate of 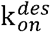 and dissociation constant 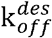, the change with time of the number of the G protein-bound as well as desensitized receptors 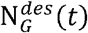 follows

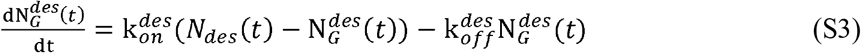

The solution of the equation is

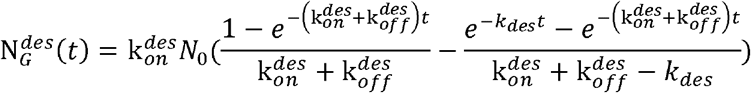

which can be re-organized to

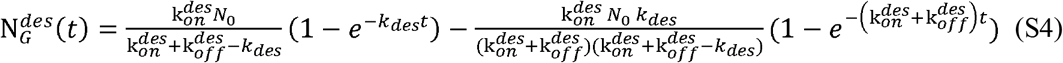

Assuming that the activated but not yet desensitized GPCRs recruit the G proteins with the association rate 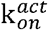 and the dissociation rate 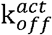, the change with time of the number of the G protein-bound but not yet desensitized receptors 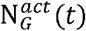 follows

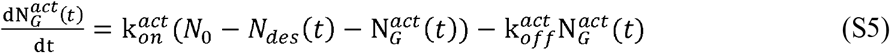

The solution of the equation is

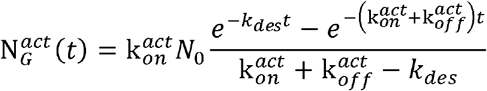

which can be re-organized to

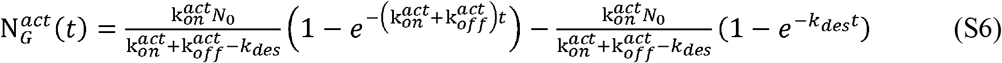

The total number of the G protein-bound receptors is

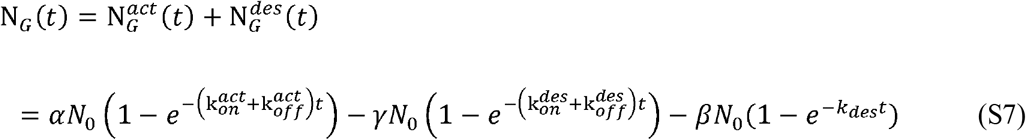

where 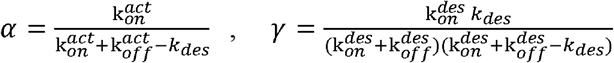 and 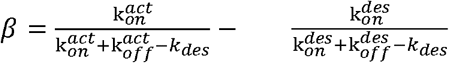

Our experimental data indicated that 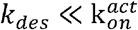 and 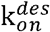. Consequently, the *γ* term can be reasobaly omitted from Eq. (S7), yielding

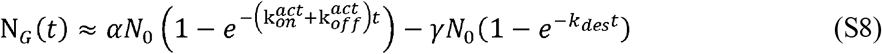

The total number of the receptors that are not occupied by the G proteins is given by

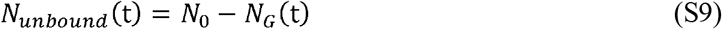

Our single-molecule measurements identified only two states in the G protein-dependent pathway: the G protein-bound state with τ_G_/τ_0_ = F_G_/F_G_ ≈ 0.85 and the G protine-unbound state with τ*_unbound_*/τ_0_ = F*_unbound_*/F_0_ ≈ 0.65. We did not see an additional effect of the desensitization by PKA on the conformation of the C-tail, which is understandable because PKA is considered to phosphorylate the intracellular loop 3. The ensemble average of the lifetime measured with FLIM is then given by

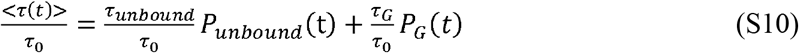

where P*_unbound_*(t) = N*_unbound_*(t)/N_0_ and P_G_(t) = N_G_(t)/N_0_.

Model Limitations. The kinetic model assumes memoryless transitions between effective states.^56^ In reality, after a G protein dissociates from the receptor, it may undergo local rebinding due to the high local concentration near the membrane, which would make the effective dissociation appear slower than the microscopic unbinding step. Furthermore, because the C-tail is a flexible polymer, the donor-to-membrane distance follows a distribution rather than a single value; the two fluorescence states we resolve likely correspond to the means of two overlapping distance distributions rather than to two sharply defined conformations.^57, 58^ The present coarse-grained model captures the dominant kinetic features of the system and reproduces the FLIM data well, but it does not exclude more complex dynamics (e.g., sub-diffusion of the G protein on the membrane, or multiple sub-states of the tail). These considerations do not invalidate our main conclusions, but they remind the reader that the extracted rates are signal-defined effective parameters that may incorporate contributions from hidden fast equilibria.

## AUTHOR INFORMATION

### Authors

**Zhao Lin -** Beijing National Laboratory for Condensed Matter Physics, Institute of Physics, Chinese Academy of Sciences, Beijing 100190, P. R.China; University of Chinese Academy of Sciences, Beijing 100049, P. R China

**Xinyao Chen -** Beijing National Laboratory for Condensed Matter Physics, Institute of Physics, Chinese Academy of Sciences, Beijing 100190, P. R.China; University of Chinese Academy of Sciences, Beijing 100049, P. R China

**Lijuan Ma -** Beijing National Laboratory for Condensed Matter Physics, Institute of Physics, Chinese Academy of Sciences, Beijing 100190, P. R.China; University of Chinese Academy of Sciences, Beijing 100049, P. R China

**Hang Fu -** Beijing National Laboratory for Condensed Matter Physics, Institute of Physics, Chinese Academy of Sciences, Beijing 100190, P. R.China; University of Chinese Academy of Sciences, Beijing 100049, P. R China

**Hao Wang-** Beijing National Laboratory for Condensed Matter Physics, Institute of Physics, Chinese Academy of Sciences, Beijing 100190, P. R.China; University of Chinese Academy of Sciences, Beijing 100049, P. R China

**Kai Yang -** School of Physical Science and Technology, Soochow University, Suzhou 215006, P. R China

## Author Contributions

## ACKNOWLEDGMENT

This work was supported by the National Natural Science Foundation of China (32471278, T2221001, 12434008, 32171228); the Chinese Academy of Sciences Project for Young Scientists in Basic Research (YSBR-104); the National Key Research and Development Program of China (2019YFA0709304).

## ABBREVIATIONS

C-tails: cytoplasmic tails
GPCR: G protein-coupled receptor
GFRET: fluorescence resonance energy transfer
BRET: the bioluminescence resonance energy transfer
H8: eighth helix
BHQ: black hole quenchers
GUV: giant unilamellar vesicles

## ASSOCIATED CONTENT

### Supporting Information Available

Additional experimental results.

## BRIEFS

This study introduces a single-fluorophore FRET approach to monitor real-time conformational dynamics of GPCR cytoplasmic tails in living cells, providing a direct molecular readout of receptor activation through distance-dependent fluorescence quenching.

## Table of Contents

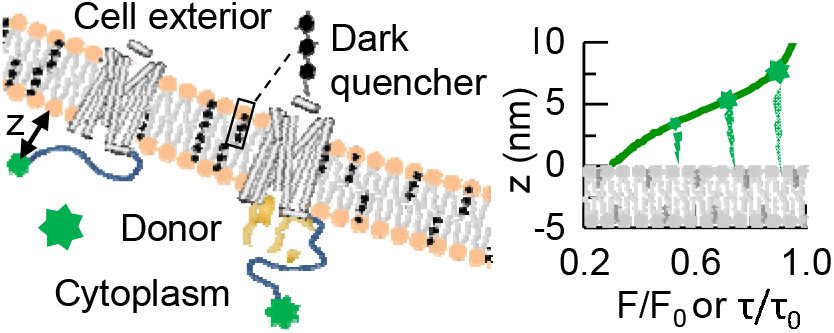

