## Supporting information for "Imaging and quantifying cytoplasmic tail dynamics of membrane receptors in living cells"

Table S1. DNA sequences used in this work.

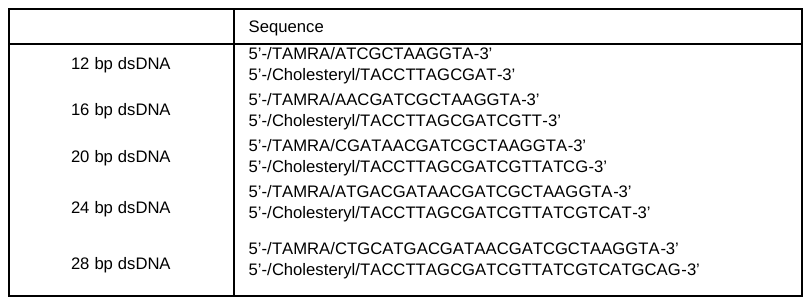

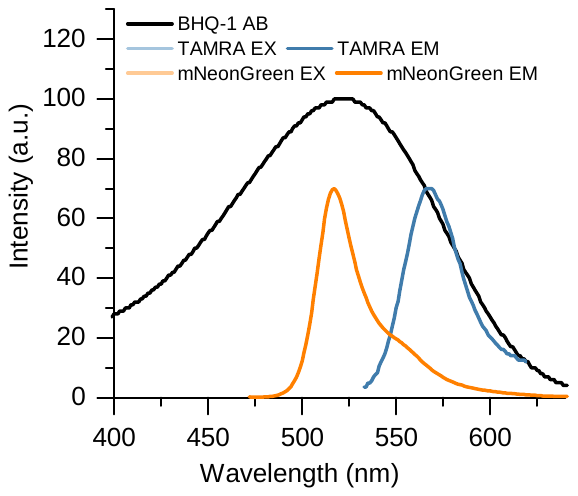

**Figure S1. Photophysical properties of the quenchers.** Comparison between the absorption spectrum of the quencher BHQ-1 and the emission spectra of TAMRA and mNeonGreen use as the donors in the current work.

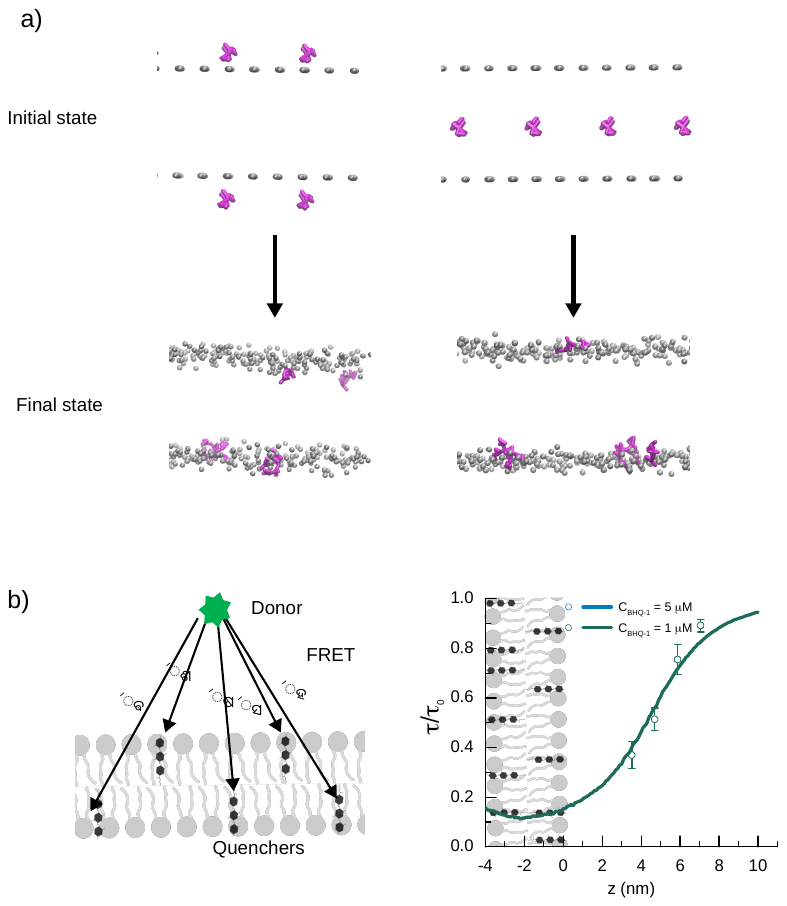

**Figure S2. Calculation of the quenching efficiency (τ/τ_0_ of F/F_0_) as a function of the donor-to-surface distance.** a) The molecular dynamics simulations of the position of the quenchers in the membrane. The BHQ-1 molecules stayed in the headgroup region (grey beads) irrespective of the initial conditions: either on the surfaces (left pannel) or in the middle plane (right pannel) of the lipid bilayer. b) The Monte Carlo simulation was applied to calculate the quenching efficiency (τ/τ_0_ of F/F_0_) as a function of the donor-to-surface distance provided the density of the quenchers according to the dsDNA calibration (Figure 1).

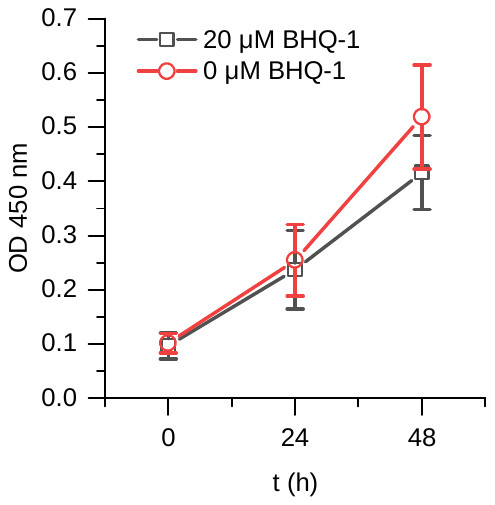

**Figure S3. Cytotoxicity of the quenchers.** The Cell Counting Kit-8 (CCK-8) assay was used to compare the viability of Hela cells in the culturing medium with (black) and without (red) BHQ-1. About 2000 cells in 200 μL medium were dispensed to a 96-well plate. After incubation for overnight, the cells were exposed to 20 μM BHQ-1 for 30 min. At 0 h, 24 h and 48h after the incubation, the CCK-8 reagent was added to the wells and incubated for 4 h. The viability of the cells was analyzed by measuring the optical density (OD) at 450 nm (1510, Thermo Fisher). Our analysis indicated that BHQ-1 does not affect the survival of the cells.

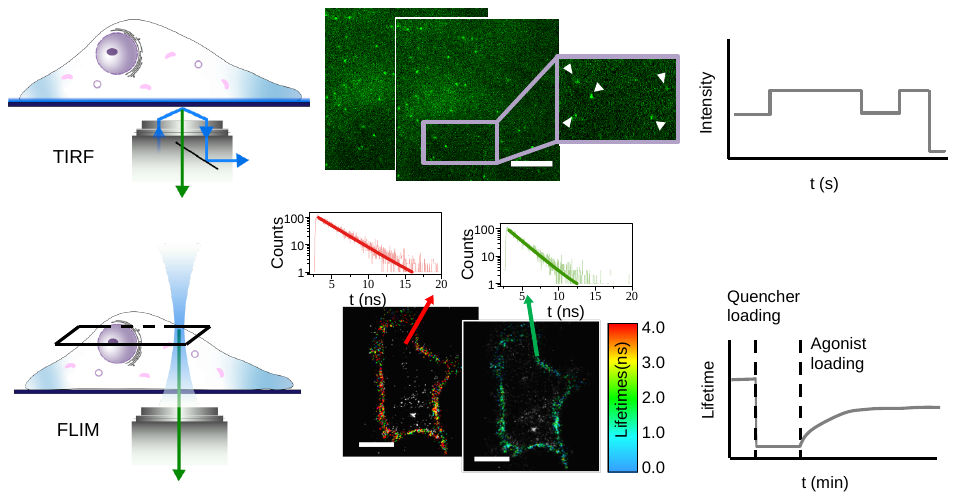

**Figure S4. Imaging platforms and data analysis.** Upper left, single-molecule imaging using TIRFM. Upper middle, representative image stacks showing fluorescent protein-labeled β_2_ARs diffusing at the cell basal plane. Upper right, fluorescent time trace of a molecule of interest. Lower left, single-cell imaging using FLIM. Lower middle, representative FLIM images at different times. Lower right, time course of lifetime averaged over a cell of interest. Scale bar: 10 μm.

**Figure S5. Demonstration of the method by measuring the dye-to-surface distance of a series of dsDNA on GUV.** a) Schematic of the GUVs platform that tethered on the bottom of the fluidic chamber. FLIM images of GUVs containing TAMRA-labeled dsDNA segments of various lengths in the presence of 5 μM BHQ-1. b) Histograms of the photon arrival times as measured by TCSPC (see Methods). Each histogram was fitted to an exponential function to obtain the corresponding lifetime. c) Statistic data of τ /τ_0_ obtained from different GUVs with dsDNA segment of various lengths. d) Comparison of the F/F_0_ (or τ/τ_0_)–distance relationship calculated by using the Monte Carlo simulation presented in Figure S2 to that measured in c). Error bars are standard derivations.

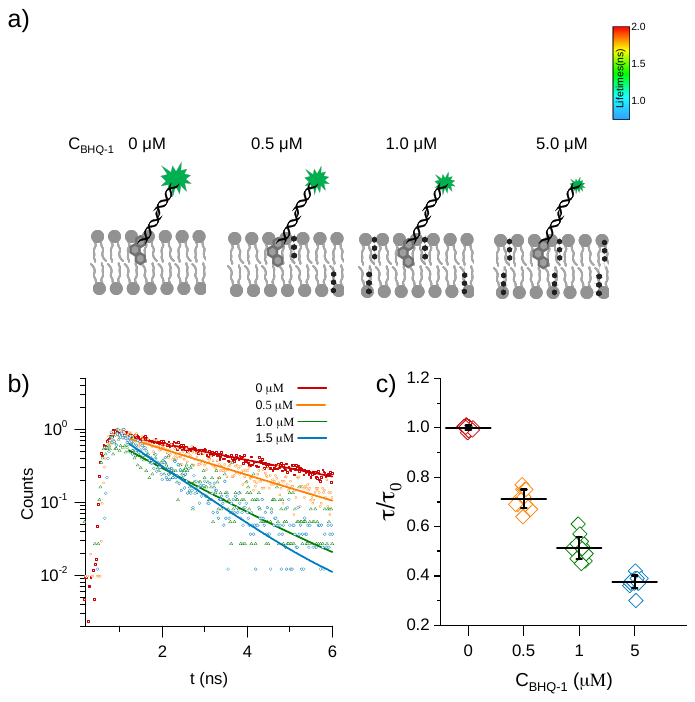

**Figure S6.** **Demonstration of the method by measuring quenching ability at various concentration of BHQ-1 on GUV.** a) Schematic and FLIM images of GUVs containing TAMRA-labeled dsDNA segments in the presence of various concentration of BHQ-1. The GUVs were tethered on the bottom of the fluidic chamber. b) Histograms of the photon arrival times as measured by TCSPC (see Methods). Each histogram was fitted to an exponential function to obtain the corresponding lifetime. c) Statistic data of τ /τ_0_ obtained from different GUVs with dsDNA segment in the presence of various concentration of BHQ-1.

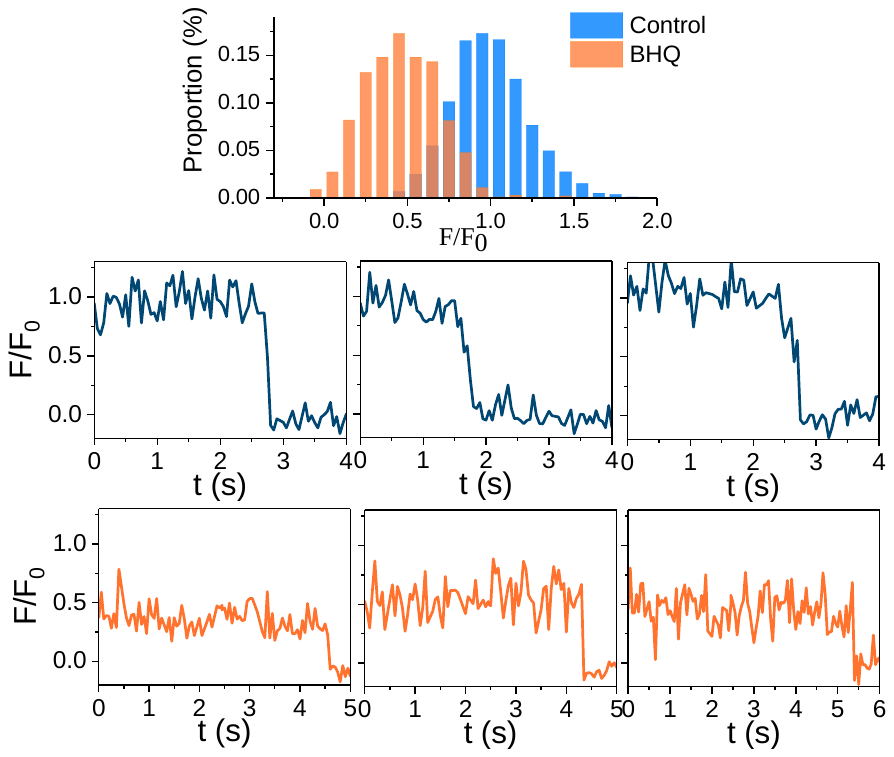

**Figure S7.** **The representative single-molecular traces and population histogram of 16 bp DNA in Hela cells before and after BHQ-1 was introduced.**

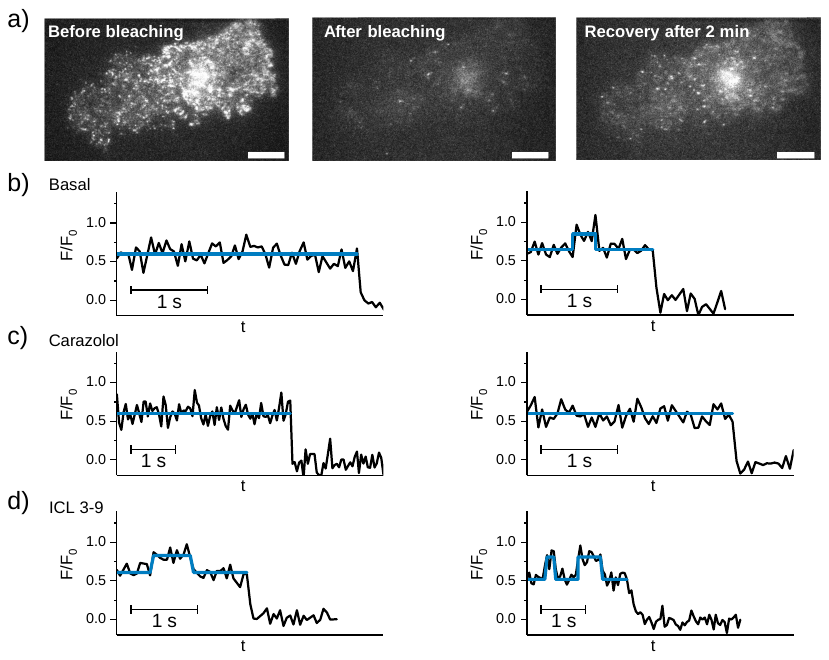

**Figure S8. TIRF imaging for single-molecule analysis.** a) TIRF images of a Hela cell expressing β_2_AR-mNG before (left), immediate after (middle) and 2 min after photobleaching. Scale bar = 10 μm. b) Representative time traces of F/F_0_ of β_2_AR**–**mNG after BHQ-1 loading. c) Representative time traces of F/F_0_ of β_2_AR**–**mNG after carazolol inhibition. d) Representative time traces of F/F_0_ of β_2_AR**–**mNG after ICL 3-9 activation. These traces show two types of molecules, including one typical stable molecule that remains in the compact state throughout the recording, and another switch molecule that alternates between the compact and extended states. We observed the increasing of stable molecules after carazolol loading, and the increasing of switch molecules after ICL 3-9 loading.

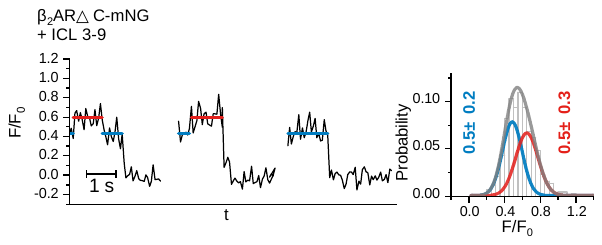

**Figure S9.** Single-molecule analysis of the β2AR△C-mNG in ICL 3-9 stimulated cells.

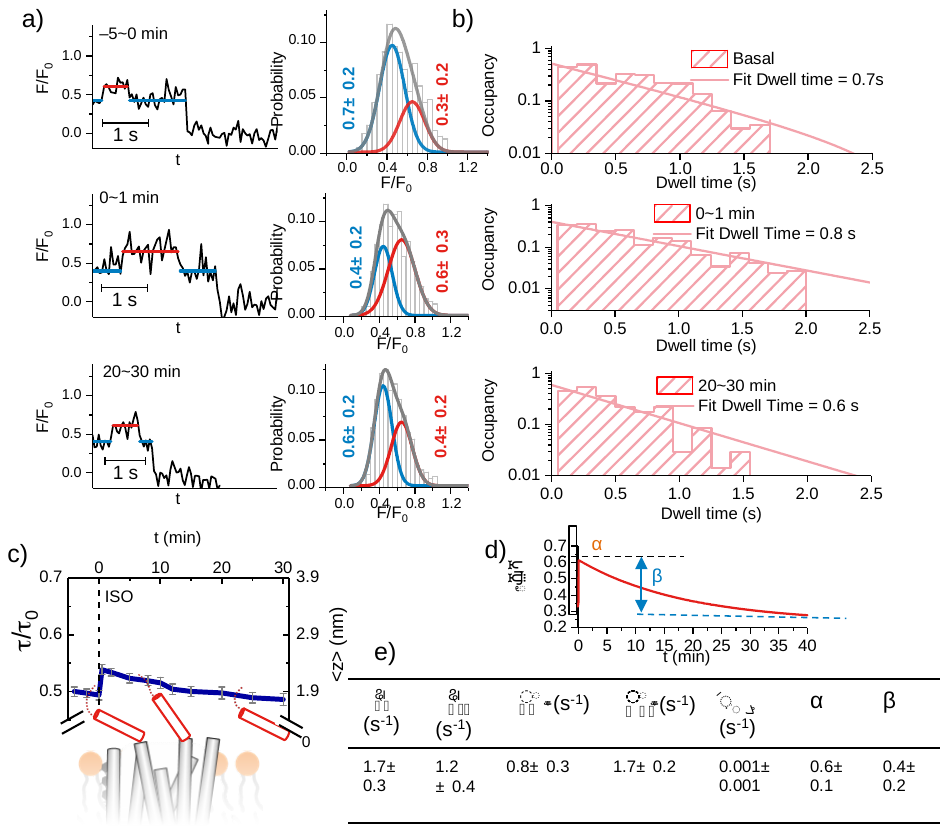

**Figure S10**. Conformational changes of the β2AR△C-mNG induced by the isoproterenol (ISO). a) Representative intensity traces collected at different time windows after the ISO loading. The histograms of the conformations are displayed on the right. b) Dwell times of the extended state in resting, immediately activated and desensitized cells. The statistics were from >100 events. c) Time course of the lifetime measured by time-lapse FLIM and the schematic of the corresponding C-terminus-to-surface distance (red cycles with error bars). The black line represents the theoretical curve derived from the kinetic model. d) The occupation probability P_G_(t) of a GPCR by a G protein using the kinetic parameters in e).
